# One gecko’s pain is another gecko’s gain: is the Moorish gecko *Tarentola mauritanica* becoming invasive in France?

**DOI:** 10.1101/2023.11.04.565611

**Authors:** Julien Renet, Théo Dokhelar, Nicolas Dubos

## Abstract

The Moorish gecko *Tarentola mauritanica* is currently expanding around the Mediterranean basin as a result of natural dispersal and anthropogenic spread. The species is observed at several sites in sympatry with other gecko species. To date, no impact has been observed on the native species and *T. mauritanica* is not considered invasive. We present an eight-year survey in southern France, where it lives in sympatry with the European leaf-toed gecko *Euleptes europaea*. The survey started when the Moorish gecko was rare which enabled us to observe an important increase in abundance. This increase was strongly correlated with a notable decline of *E. europaea*, explaining 49% of transect-specific temporal variation in abundance. We suspect that the increase in *T. mauritanica* density is causally related to this decline and recommend intensive monitoring of the species throughout the Mediterranean Basin to determine whether or not the species should be classified as invasive.

## Introduction

Global changes (e.g. international trade, climate change) have caused shifts in the distribution of biodiversity, especially in squamates (Pyšek et al., 2010; Moreno-Rueda, 2012; Ceia-Hasse et al., 2014; Bonino et al., 2015). This process leads or will lead to the contraction of many species ranges and the colonisation of new geographical areas for some species (Nasrabadi et al., 2018; Gómez-Cruz et al., 2021).

Native to North Africa and Iberian Peninsula, the Moorish gecko *Tarentola mauritanica,* currently expanding in Southern Europe, appears to be particularly responsive to globalisation. The colonisation patterns of the species are probably a result of a combination of anthropogenically mediated-spread and natural dispersal (Harris et al., 2004a, b; Perera & Harris, 2008; Rato et al., 2010, 2012; Silva-Rocha et al., 2022) thanks in part to the resilience of an Iberian core during the Pleistocene (Rato et al., 2010). It is also possible that the natural spread of the species is facilitated by climate change, since the species is present in warmer and more arid regions (Rato & Carretero, 2015; Rato et al., 2015).

The discovery of *T. mauritanica* in new localities is often attributed to recent accidental introductions by humans, particularly on islands (e.g. Jesus et al., 2008; Barreiros et al., 2010; Mačát et al., 2014; Deso et al., 2020; Rato et al., 2021; Strachinis et al., 2023). The species has become established in very distant areas from its native range, such as Mexico (Ortiz-Medina et al., 2019), Argentina (Baldo et al., 2008; Díaz-Fernández et al., 2019), Uruguay (Baldo et al., 2008), California (Mahrdt, 1998) and Florida (Rochford & Krysko, 2019). Within the Mediterranean Basin, the species recently extended its distribution northward, where several colonisation fronts were identified (Fig. 1). One front is located in Southwestern France, along a major terrestrial transportation route (Garonne valley), already identified as a factor of spread in the past for several other Mediterranean species (Berroneau, 2014). Another one is located in the Rhône valley where *T. mauritanica* has reached the Isère French department (Grossi & Fonters, 2015). A northward spread was also recorded in the Durance valley and in the Var valley (Alpes-Maritimes) (Renet J unpublished data).

**Figure 1.**
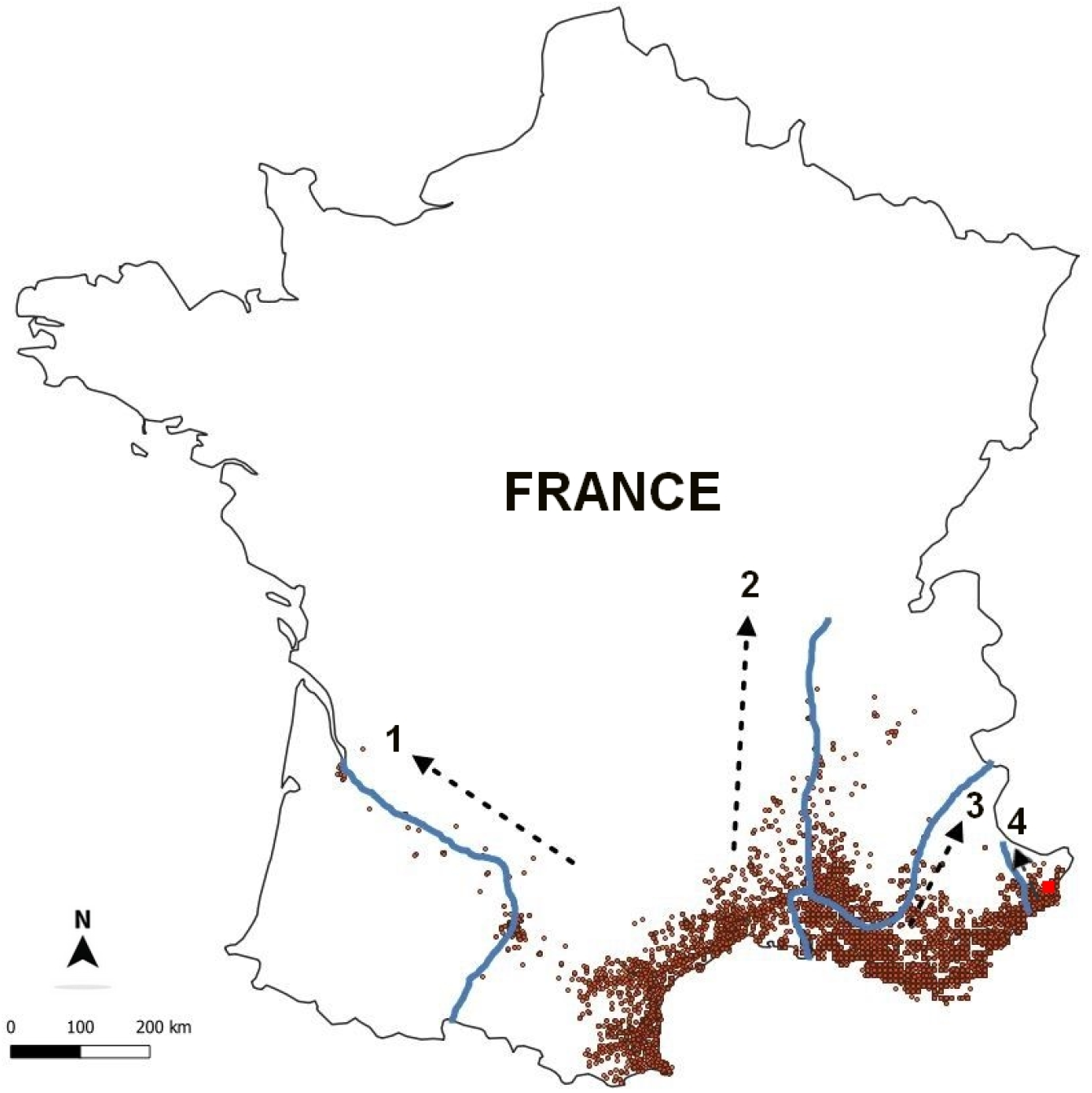
Distribution of the Moorish gecko *Tarentola mauritanica* in mainland France. Orange points correspond to species occurrence. Black arrows correspond to the direction of its spread in the Garonne (1), Rhône (2), Durance (3) and Var (4) valleys. The red square shows the location of the study site in the Alpes-Maritimes. Data OpenObs (INPN – SINP) Leaflet – Map data © OpenStreetMap, imagery © CartoDB.

Whether *T. mauritanica* has an impact on native species is currently subject to debate from a conservation point of view. For example, the decline of the Common wall lizard *Podarcis muralis* in urban environments was attributed to the Moorish gecko’s arrival, although no scientific evidence has been reported yet (Geniez & Cheylan, 2012). Conversely, Simbula et al. (2019) highlighted that co-existence with *P. muralis* (and other species) does not lead to competitive interactions but rather to a strong partitioning of ecological niches likely due to extensive separation in microhabitat (e.g. vertical surfaces) and the period of activity of the two species (i.e. catemeral vs. diurnal). Despite their activity being asynchronous, both species feed on terrestrial arthropods, which might to reduce local resources. Additionally, *P. muralis* may share retreat sites for thermoregulation with *T. mauritanica*, which suggests possible competition for micro-habitats.

In the Western Mediterranean, *T. mauritanica* co-occurs with two other species of nocturnal geckos, the Mediterranean gecko *Hemidactylus turcicus* and the European leaf-toed gecko *Euleptes europaea*. The latter is narrowly distributed and it mainly occupies islands, which makes it a species of high conservation value (Salvidio et al., 2010). It is classified as Near-Threatened at the global level (Temple & Cox, 2009) and in danger of extinction in Southeastern France (Marchand et al., 2017).

Several cases of co-occurrence between *T. mauritanica* and *E. europaea* have been documented throughout the natural range of *E. europaea*, notably on Corsica, Sardinia and some satellite islands (e.g. Girraglia, Finnocchiarola, etc.), on islands in Tuscany (e.g. Elba, Montecristo, etc.) (Delaugerre & Cheylan, 1992; Delaugerre & Guyot, 1995), in Provence (Ratonneau, Levant Island, Port-Cros, Porquerolles) (Delaugerre, 1981; Astruc et al., 2009; Deso et al., 2018; Deso et al., 2020) and on several mainland localities in the Alpes-Maritimes, Liguria and Tuscany (Renet et al., 2008; Salvidio et al, 2010; Radi & Zuffi, 2022). *Tarentola mauritanica* is suspected to have caused local extinctions in *E. europaea*. This is the case in the island of Bendor, which is now occupied solely by *T. mauritanica*, although the island was home to a *E. europaea* population, before *T. mauritanica* arrived (Mourgue, 1910; Jahandiez et al., 1933; Ineich et al., 2019). The presence of *T. mauritanica* was also suspected to cause the extirpation of *E. europaea* on the largest island of the Galite archipelago (Tunisia), Galita. Indeed, the island is largely occupied by *T. mauritanica*, while no *E. europaea* has never been observed despite several sampling campaigns (Lanza & Bruzzone, 1959; Abbes et al., 2008). This is supported by its presence on smaller peripheral islands which are free of *T. mauritanica* (Delaugerre et al., 2011; Corti et al., 2022). A similar situation has been observed on the Lérins archipelago (Cannes, France) where *T. mauritanica* largely occupies the two largest islands (i.e. Sainte-Marguerite and Saint-Honorat) while *E. europaea* is confined to adjacent islets (i.e. Saint-Féréol and La Tradelière) (Renet & Martinerie, 2011; Renet et al., 2013). In spite of this, no monitoring protocol has been set to study in detail the consequences of the co-existence of these two species.

This study presents an 8-year monitoring of a population of *E. europaea* at a site where it is found in sympatry with *T. mauritanica*, with both species being recorded during application of a standard transect protocol. We provide evidence for the collapse of *E. europaea* and the rise of *T. mauritanica*. The aim of this study is to quantify population trends for both species, discuss the causes that could explain this observation and to provide guidelines for further research and management.

## Material & Methods

### Study site and data collection

The sampling site is located in a group of south-oriented limestone ledges in the Alpes-Maritimes, at 610 m a.s.l. on the communes of Eze and La Turbie. The site corresponds to the thermo-Mediterranean floristic stage (Quézel & Médail, 2003), with a shrub layer mainly composed of *Pistacia lentiscus* L., *Rhamnus alaternus* L., *Quercus ilex* L., and *Euphorbia dendroides* L. A strategic military track was integrated into the site by the end of the 19^th^ century, which facilitated access and sampling campaigns from one year to the next. The track consists of structurally homogeneous rocky walls on one side and a partially jointed dry-stone wall about 80 cm high on the other side.

We recorded the number of reptiles along four geolocated 110 m-long transects, set up on the military track at least 100 m apart from each other. These were surveyed between 2009 and 2017 for 40 minutes each, systematically by two experienced observers on two to three occasions in spring (April to June) and autumn (September to October) for a total of 40 sessions. We recorded *E. europaea* and *T. mauritanica* at night when weather conditions were appropriate (i.e., in the absence of precipitation and wind, with temperature ranging between 12 and 22°C) using headlamps (500 lumens). The start times of the surveys varied from 20:00 to 21:30 depending on the season, and did not go beyond 00:45. We carefully inspected micro-habitats on both sides of the track. Each observer was assigned to only one edge of the runway to avoid double counting. Only sexually mature adults were counted. We did not count juveniles, because they are more difficult to detect due to their smaller size.

### Statistical analysis

#### Assessing temporal trends

We fitted Poisson Generalised Linear Mixed models (GLMMs) to the count data using the lme4 package (Bates et al., 2015) in R v.4.2.0 (R Core Team, 2021) (this R package was used for all analyses here). This allowed us to test for of linear temporal trends in the number of observations of the two sympatric species *E. europaea* and *T. mauritanica*. We built a null model which included three adjustment variables: We accounted for differences in abundance between species using a ‘species’ fixed effect, while the transect number and the sampling session were treated as random effects. We build one model assuming a common temporal trend among both species (i.e., with year as an additive continuous effect). We then assessed a model assuming a different temporal trend between species using season as a fixed effect and an interaction term between species and year. We assessed these models using the dredge function from the *MuMIn* package (Barton, 2020) on the basis of information theory (second order Akaike Information Criterion, AICc ; Burnham & Anderson, 2002). We chose the model with the best trade-off between goodness of fit and the number of variables included, i.e. with the lowest AICc (Wagenmakers & Farrell, 2004; Bolker, 2008; Bolker, 2021). Models within a delta AICc of 2 of the best model were considered to have substantial statistical support and given further consideration (Burnham and Anderson, 2002). We computed the Marginal R² (Rm²), estimating the variance explained by the fixed factors, and conditional R² (Rc²), estimating the variance explained by both the fixed and random factors (Nakagawa et al., 2017) using the R package *performance* (Lüdecke et al., 2021).

We performed pairwise post-hoc tests with the *emmeans* function from the *emmeans* package (Lenth, 2021) in order to test the difference in the number of observations between the two species through the eight sampling years. All graphics were generated using the R package *ggplot2* (Wickham, 2016).

#### Assessing non-linear temporal trends

We tested the robustness of our linear assumptions on temporal trends using Generalised Additive Mixed Models (GAMMs) with mgcv package v. 1.8-42 (Wood & Scheipl, 2014). GAMMs fit the data with optimal degrees of freedom and are appropriate to describe non-linear temporal trends or fluctuations. We built one model for each species separately, with species count as response variable and year with a spline effect as predictor. We included transect and sampling session as random effects.

#### Assessing the effect of T. mauritanica on E. europaea

We quantified the effect of *T. mauritanica* abundance on *E. europaea*. We used GLMM assuming a Poisson distribution with *E. europaea* counts as response variable and *T. mauritanica* counts as predictor (hereafter, *m1*). We inspected for differences in abundance between transects and sampling sessions using random effects. We took into account the temporal dependence between consecutive years in each transect by including a ’year’ random intercept and slope effect, which we allowed to vary among transects with a random intercept term. We compared the AICc of the model with that of a null model (which includes adjustment variables only; hereafter *m0*) to verify that the inclusion of *T. mauritanica* counts is informative. Finally, we quantified the amount of temporal variance specific to each transect explained by *T. mauritanica* abundance. To do so, we computed the ratio between the between-year variance of *m0* and the between-year variance of *m1* estimated with the random effects.

## Results

Between 2009 and 2017 *E. europaea* observations decreased by 78.49 % while *T. mauritanica* observations increased by 7329 %. Based on AICc scores, we found strong statistical support for species-specific temporal trends but no effect of season on species observation (Rm² = 0.886, Rc² = 0.965; Table 1). Pairwise emmeans post-hoc test showed that from 2009 to 2014 *E. europaea* were significantly more abundant than *T. mauritanica* (Table 2, Fig 2) but that in 2015 and 2017 the opposite trend was observed (Table 2, Fig 2). The GAMMs showed that the trends were non-linear (*E. europaea*: Estimated degrees of freedom = 2.85; P > 0.0001; *T. mauritanica*: Estimated degrees of freedom = 2.74; P > 0.0001; Fig 3). The abundance of *E. europaea* started to decline slowly near 2011 and declined linearly between 2012 and 2017. *T. mauritanica* showed a rapid linear increase between 2009 and 2012 approximately, which was less pronounced between 2012 and 2017. We found strong statistical support for a negative impact of *T. mauritanica* on the abundance of *E. europaea* (Estimate ± SE = -0.06 ± 0.005, *P* < 0.00001; ΔAICc (*m0*-*m1*) = 28.5). The abundance of *T. mauritanica* explained 49% of the transect-specific inter-annual variance.

**Table 1.** Model ranking for the effect of species and year on gecko count (*E. europaea* and *T. mauritanica*). Marginal (Rm²) and conditional (Rc²). R² values indicate the goodness-of-fit of the models.

| Explanatory variable | AICc | dAICc | df | $R_m^2$ | $R_c^2$ |
| --- | --- | --- | --- | --- | --- |
| Species*Year | 1611.2 | 0.00 | 18 | 0.886 | 0.965 |
| Season + Species*Year | 1612.9 | 1.75 | 19 | 0.886 | 0.965 |
| Species+Year | 3182.1 | 1570.88 | 11 | 0.536 | 0.858 |
| Season + Species + Year | 3183.7 | 1572.53 | 12 | 0.536 | 0.857 |
| Species | 3190.1 | 1578.87 | 4 | 0.531 | 0.856 |
| Season + Species | 3191.9 | 1580.75 | 5 | 0.527 | 0.857 |
| Season | 4261.8 | 2650.63 | 4 | 0.026 | 0.700 |

**Table 2.** Summary of the pairwise emmeans post hoc test using *Euleptes europaea* and *Tarentola mauritanica* as contrast.

| Year | Estimate | SE | df | z.ratio | p.value |
| --- | --- | --- | --- | --- | --- |
| 2009 | 102.286 | 38.8489 | 302 | 12.185 | <0.0001 |
| 2010 | 44.538 | 12.4907 | 302 | 13.537 | <0.0001 |
| 2011 | 8.776 | 1.0625 | 302 | 17.941 | <0.0001 |
| 2012 | 5.129 | 0.5814 | 302 | 14.423 | <0.0001 |
| 2013 | 2.075 | 0.1921 | 302 | 7.888 | <0.0001 |
| 2014 | 1.198 | 0.1089 | 302 | 1.989 | 0.0476 |
| 2015 | 0.562 | 0.0515 | 302 | -6.290 | <0.0001 |
| 2017 | 0.296 | 0.0384 | 302 | -9.379 | <0.0001 |

**Figure 2.**
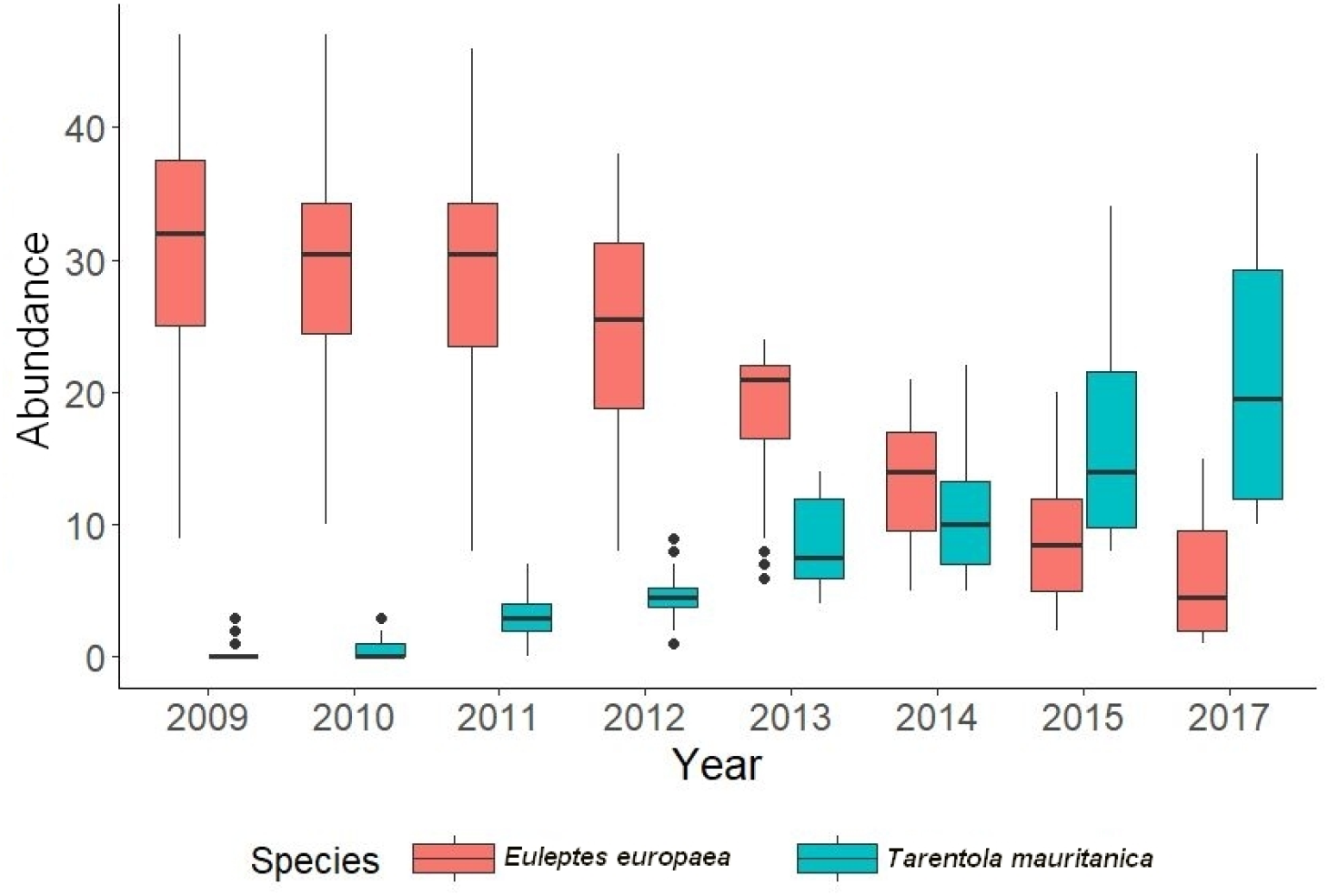
Abundance of *Euleptes europaea* and *Tarentola mauritanica* along four transects throughout eight sampling years. Boxes indicate the 25%, 50% (median) and 75% quartiles. Lines show minimum and maximum values excluding outliers (indicated as dots).

**Figure 3.**
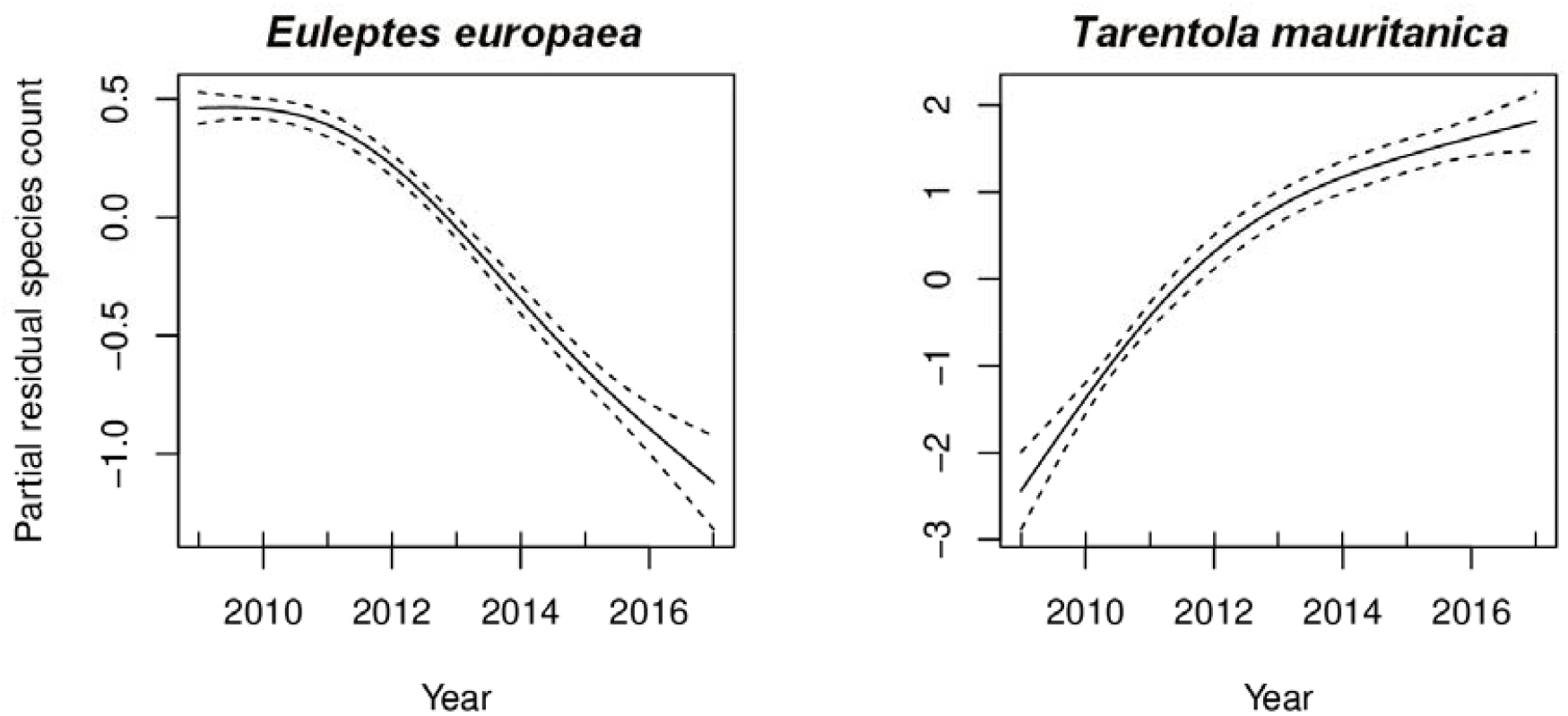
Temporal variation in abundance of *Euleptes europaea* (left panel) and *Tarentola mauritanica* (right panel). Curves represent fitted values from GAMM, with year as a spline effect. Dotted lines represent 95% confidence intervals.

## Discussion

We provide evidence of a decline in the abundance of *E. europaea* which is worryingly correlated with a very large increase in the number of *T. mauritanica* observed within the same microhabitats. We believe that this may be linked to access to refuges. The small size of *E. europaea* allows it to select narrower crevices that are difficult for adult *T. mauritanica* to access. On the other hand, these narrow fissures could be accessible to immature *T. mauritanica*. At present, our data do not allow us to carry out a detailed analysis of habitat selection by these two species.

The cause of the increase in *T. mauritanica* is currently unknown. The species was already considered common in Southeastern France at the beginning of the 19^th^ century (de Serre, 1822; Caziot, 1922) and no habitat change was recorded at the study site before or during the monitoring period. In addition, unrestrained urbanisation is likely to have facilitated the spread of individuals translocated from other areas (e.g., through the transportation of construction materials or ornamental plants). This may have led to increased interspecific competition, especially for access to food resources and refuges in rocky habitats (crevices). *Tarentola mauritanica* is characterised by a strong ecophysiological plasticity (Carretero, 2008; Rato & Carretero, 2015), an efficient mode of reproduction (i.e., continuous spermatogenesis: Angelini et al., 1983; Picariello et al., 1989) likely to vary with climate change (e.g. possible extension of the ovarian seasonal cycle length) (Clarke & Zani, 2012), strong territoriality and frequent agonistic behaviours (Lisičić et al, 2012; Salvador, 2016). Although both nocturnal and diurnal, *T. mauritanica* is mainly nocturnal (Lisičić et al, 2012), hence potential overlaps with the activity of *E. europaea*. Overall, these traits suggest a high potential of invasiveness. Further research that focuses on determination of the climate niche of the species may therefore improve our understanding of climate change effects and its spread. Notwithstanding the high potential invasiveness of *T. mauritanica*, its impact may be mitigated in the case of niche partitioning.

Evidence of strong spatial segregation between *T. mauritanica* and *H. turcicus* was previously found without proving the existence of a mutual negative impact (Lisičić et al., 2012). Competition for spatial niches leading to habitat partitioning may occur at our study site. Olfactory marking induced by the presence of a potential competitor or predator can generate physiological stress that leads to different behavioural responses (Febrer-Serra et al., 2023). In particular, in the presence of a nocturnal co-occurring species (i.e. the Black rat), *E. europaea* tends to remain sheltered in crevices (Delaugerre et al., 2019). This may have affected our ability to detect the species and led to a large underestimation of the number of individuals present. Nevertheless, in Cala Violina on the Tuscan coast, *E. europaea* did not show any atypical spatial behaviour (e.g. emerge from crevices less frequently) despite the presence of *T. mauritanica* in similar densities (Radi & Zuffi, 2022).

The question of high predation pressure also arises, but studies on the trophic ecology of *T. mauritanica* indicate that it feeds almost exclusively on arthropods (Gil & Perez-Mellado, 1994; Hódar et al., 2006; Martins et al., 2022). However, some cases of predation on several species of Lacertidae (Salvador 1978; Franco 1980; Pellitteri-Rosa et al., 2015) have been documented in other areas. For example, the DNA of the wall lizard *Teira dugesii* was detected in nearly 27% of *T. mauritanica* samples from Madeira Island (Martins et al., 2022). Predation of *H. turcicus by T. mauritanica* has also been observed, (Rieppel 1981; Gonzalès de la Vega 1988; Bauer 1990). Given the large difference in body size between *T. mauritanica* (total length 15 cm) and *E. europaea* (total length 8 cm), predation may occur occasionally on juvenile or adult *E. europaea*, but this alone cannot explain the extent of the observed decline. Apart from potential predation by *T. mauritanica*, competition for food resources cannot be ruled out, given the high similarity in trophic ecology between the two species (Oneto et al., 2008; Hódar et al., 2006; Martins et al., 2022).

Finally, the possibility of parasite transmission cannot be excluded. *Tarentola mauritanica* often carry intestinal parasites (Helminth) (El-Rghibi et al., 2022), hemoparasites (*Leishmania tarentolae*, haemogregarines of the Genera *Hepatozoon* ssp.) (Rioux et al., 1969; Tomé et al., 2016; Parejo-Pulido et al., 2023) and acari (geckobians) (Girot, 1968; Bertrand et al., 2012) which may impact the physiology, the immunity and the behaviour of their hosts (Oppliger et al., 1996; Oppliger & Clobert, 1997; Amo et al., 2005). The introduction of non-indigenous parasites transmitted to the local *T. mauritanica* population by fortuitous individual displacement is also to be considered, along with their potential impact on physiology and population dynamics.

Our study does not allow us to certify that the increase in the occurrence of *T. mauritanica* is causally related to the decline of *E. europaea*. Nevertheless, the strong correlation between the collapse of the population of *E. europaea* and the increase in the number of *T. mauritanica* raises questions regarding the conservation of *E. europaea*. *Tarentola mauritanica* was introduced very recently on two islands in Provence (Port-Cros, Ile du Levant) (Deso et al., 2021) which could provide a more rigorous test of a direct negative impact by *T. mauritanica* on *E. europaea* populations. Most of the islands of Provence are home to high-density *E. europaea* populations and benefit from a high level of protection (Port-Cros National Park and Calanques National Park). We encourage the urgent implementation of biocontrol strategy and robust monitoring at our study site and more widely at sites where *T. mauritanica* co-occurs with other gecko species beyond its native range, especially on islands. This will avoid the entry of new specimens of *T. mauritanica* and improve the understanding of the colonisation dynamics of *T. mauritanica* and its biotic interactions. In the meanwhile, the invasive status of *T. mauritanica* remains an open question.

## Acknowledgments

We sincerely thank Antoine Renet for his participation in the nocturnal monitoring of geckos. We also thank Eric Durand (Naturalia Environnement) and Andrea Villa (Institut Català de Paleontologia Miquel Crusafont) for the constructive discussions on this issue. Three reviewers helped to improve the quality of this manuscript, and we thank them warmly.

